# Distribution of calbindin-positive neurons across areas and layers of the marmoset cerebral cortex

**DOI:** 10.1101/2024.01.05.574341

**Authors:** Nafiseh Atapour, Marcello G. P. Rosa, Shi Bai, Sylwia Bednarek, Agata Kulesza, Gabriela Saworska, Katrina H. Worthy, Piotr Majka

## Abstract

The calcium-binding protein calbindin is selectively expressed in specific neuronal populations of the cerebral cortex, including major classes of inhibitory interneurons. We have charted the distribution of calbindin-positive (CB^+^) neurons across areas and layers of the entire marmoset cortex using a combination of immunohistochemistry, AI-based image segmentation, 3-dimensional reconstruction, and cytoarchitecture-aware registration. CB^+^ neurons formed 10-20% of the cortical neuronal population, occurring in higher proportions in areas corresponding to low hierarchical levels of processing, such as sensory cortices. Although CB^+^ neurons concentrated in the supragranular layers, there were clear trends in laminar distribution: the relative density in infragranular layers increased with hierarchical level, and the density in layer 4 was lowest in areas involved in sensorimotor integration and action planning. These results reveal new aspects of the cytoarchitectural organization of the primate cortex and demonstrate an efficient approach to mapping the full distribution of neurochemically distinct cell types throughout the brain, readily applicable to most mammalian species and parts of the nervous system.

## Introduction

Precise data on the distribution and prevalence of different neuronal types is a key to the creation of biologically realistic models that can help us understand the emergence of complex patterns of brain activity (Markram et al., 2015; Wang, 2020). One of the essential criteria for characterizing neuronal classes is the cellular expression of molecules such as neurotransmitters, neuromodulatory peptides, and calcium-binding proteins. This type of characterization is often used to distinguish classes of cells that prove, upon additional work, to have distinct functional features and anatomical connections (Markram et al., 2004, De Felipe et al., 2013; Huang and Paul, 2019).

Areas of the cerebral cortex vary in terms of proportions of neurons belonging to different classes and their distributions across layers (e.g. Dombrowski et al., 2001; Scala et al., 2019). However, much research is still needed to fully understand the extent to which areas differ in terms of cellular content, and to form a comprehensive set of rules to explain this variety. For example, to date, comprehensive spatial maps of classes of neurons defined by expression of calcium-binding proteins have only been obtained in the mouse brain (Kim et al., 2017; Bjerke et al., 2020). In comparison, our knowledge of this subject in the primate cortex remains less complete, despite recent progress in mapping the distribution of cells expressing different molecules based on transcriptomics (Chen et al., 2023; Krienen et al., 2023) and receptor radioautography (Froudist-Walsh et al., 2023). A comprehensive knowledge of the distribution of different neuronal types in non-human primates is particularly important given the marked elaboration of the cortex in primate evolution and their essential role in translational research (Mitchell et al., 2018; Lear et al., 2022).

Here we report on the full distribution of neurons expressing the calcium-binding protein calbindin-D28K (CB^+^ neurons) in the cortex of a non-human primate (marmoset monkey; *Callithrix jacchus*). Calbindin (CB) can increase the neuronal calcium buffering capacity, which previous studies have demonstrated, leads to a modulatory effect on synaptic plasticity, memory, and other facets of behavior (Chard et al., 1995; Harris et al., 2016; Li et al., 2017; Molinari et al., 1996). In addition, it has been suggested that intracellular CB has protective roles which are important in the context of the development of neurodegenerative disorders and recovery from brain injury (German et al., 1992; Hof and Morrison, 1991; Yamada et al., 1990; Atapour et al., 2022). CB^+^ neurons in the primate cortex are, in vast majority, inhibitory (GABAergic) interneurons (Hendry et al., 1989; Condé et al., 1994; De Felipe 1997; De Felipe et al., 1999; Bourne et al., 2007), but also include, in some areas, a smaller and more variable population of pyramidal cells (Kondo et al., 1999). The CB marker gene (CALB1) is mainly expressed by the somatostatin (SST) and lysosome-associated membrane protein 5 (LAMP5) subclasses of GABAergic cells, which are among the five main categories of interneurons in the mammalian brain (Hodge et al., 2019; Bakken et al., 2021; Yao et al., 2021; Chen et al., 2023).

Previous studies have described the distribution of CB^+^ neurons, using either qualitative or quantitative (stereology-based) methods, in specific regions of the primate cortex such as prefrontal (Dombrowski et al., 2001), temporal (De Felipe et al., 1999; Kondo et al., 1999), auditory (Morino-Wannier et al., 1992) and visual (Goodchild and Martin 1998; Bourne et al., 2007) areas. However, we are still lacking a comprehensive spatial analysis of the distribution of CB^+^ neurons across the entire cortex of any primate. In the present study, the organizational principles governing the distribution of CB^+^ neurons across areas and layers of the marmoset cortex were defined using a computational workflow that incorporates artificial intelligence-based identification of neuronal bodies, 3-dimensional reconstruction, and cytoarchitecture-aware registration (Majka et al., 2021), combined with validation by classical stereology.

The marmoset is one of the species for which comprehensive data on the areal and laminar distributions of cortical neurons are available (Atapour et al., 2019), and it has been the subject of large-scale studies focused on structural and functional connectivity (Liu et al., 2020; Majka et al., 2020; Theodoni et al., 2021; Schaeffer et al., 2022; Tian et al., 2022). To allow further studies, the present datasets are also provided for download in a spatially oriented, three-dimensional form (NIFTI files, available as Supplemental materials to this paper and at https://www.marmosetbrain.org/whole_brain_cb_maps).

## Results

We have applied immunohistochemical techniques to reveal the locations of CB^+^ neurons (Fig. 1A) in the brains of 3 young adult marmosets (one cerebral hemisphere each). From the resulting coronal sections in one of the cases (CJ1741), we selected radial strips encompassing all cortical layers from each of the currently recognized cytoarchitectural areas of the cortex in this species (e.g. Fig. 1B, C). Manual annotation of the positions of neurons in these strips (Fig. 1D, E), using the workflow described by Atapour et al. (2019), resulted in a library of 4,072 counting boxes, which provided the basis for training a U-Net architecture Convolutional Neural Network (U-Net CNN, Ronneberger et al., 2015; Xie et al., 2018) and its evaluation (Fig. 2A, B). We found that the U-Net CNN-generated neuronal densities reflect a consensus between the estimates obtained by different human experts (Fig. 2D, E; Supplemental Figure S1). The U-Net CNN was then used to compute the densities of CB^+^ neurons in every fifth section across the entire cortex (e.g. Fig. 2F, G). Adjacent sections stained for Nissl substance or NeuN (Neuronal nuclear protein; a neuron-specific marker; Mullen et al., 1992), myelin and cytochrome oxidase were used to identify cortical areas and layers, as detailed in the *STAR* ★ *METHODS* section. This allowed the evaluation of the relations between the density of CB^+^ neurons and categories of areas defined by anatomical, functional, (hierarchical levels; Markov et al., 2014; Theodoni et al., 2021) and structural (type of lamination; Dombrowski et al., 2001; John et al., 2022) measures.

**Figure 1.**
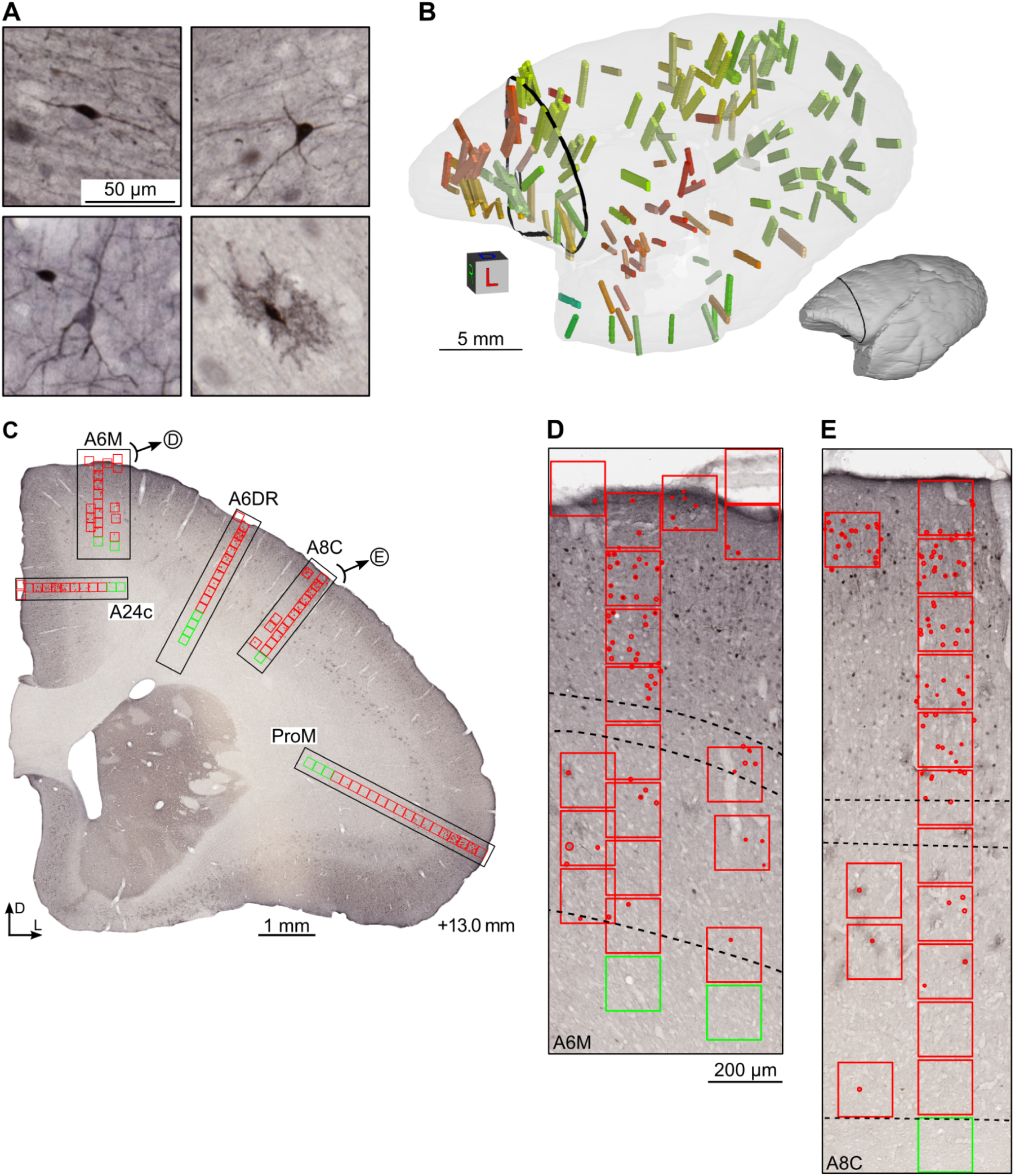
Manual annotation of calbindin-positive (CB^+^) neurons across the cerebral cortex. (A) Representative types of CB^+^ neurons (from top left to bottom right): bipolar, multipolar, small pyramidal, and neurogliaform neurons. (B) Locations of 163 image strips sampled in case CJ1741, visualized against a semi-transparent rostrolateral view of the 3D reconstructed left hemisphere (opaque model presented as a thumbnail for clarity). Different colors of the image strips correspond to the areas they were derived from (see e.g. Supplemental Figure S2D), and the black contour indicates the coronal level of the section presented in panel C. (C) Coronal cross-section taken approximately at the interaural +13.0 mm (Paxinos et al. 2012) containing five image strips (black rectangles) from areas A24c, A6M, A6DR, A8C, ProM, clockwise. (D, E): Strips from areas 6M and 8C shown at high magnification. Counting boxes are represented with red (gray matter) or green (white matter) squares of 150 µm in size. The thin, dashed band in the middle of the strips indicates the boundaries of layer 4, while the bottom dashed line shows the border between layer 6 and the white matter. The laminar boundaries were determined by comparison with adjacent Nissl-stained sections. The complete set of image strips is provided as supplemental materials and available for download from https://www.marmosetbrain.org/whole_brain_cb_maps.

**Figure 2.**
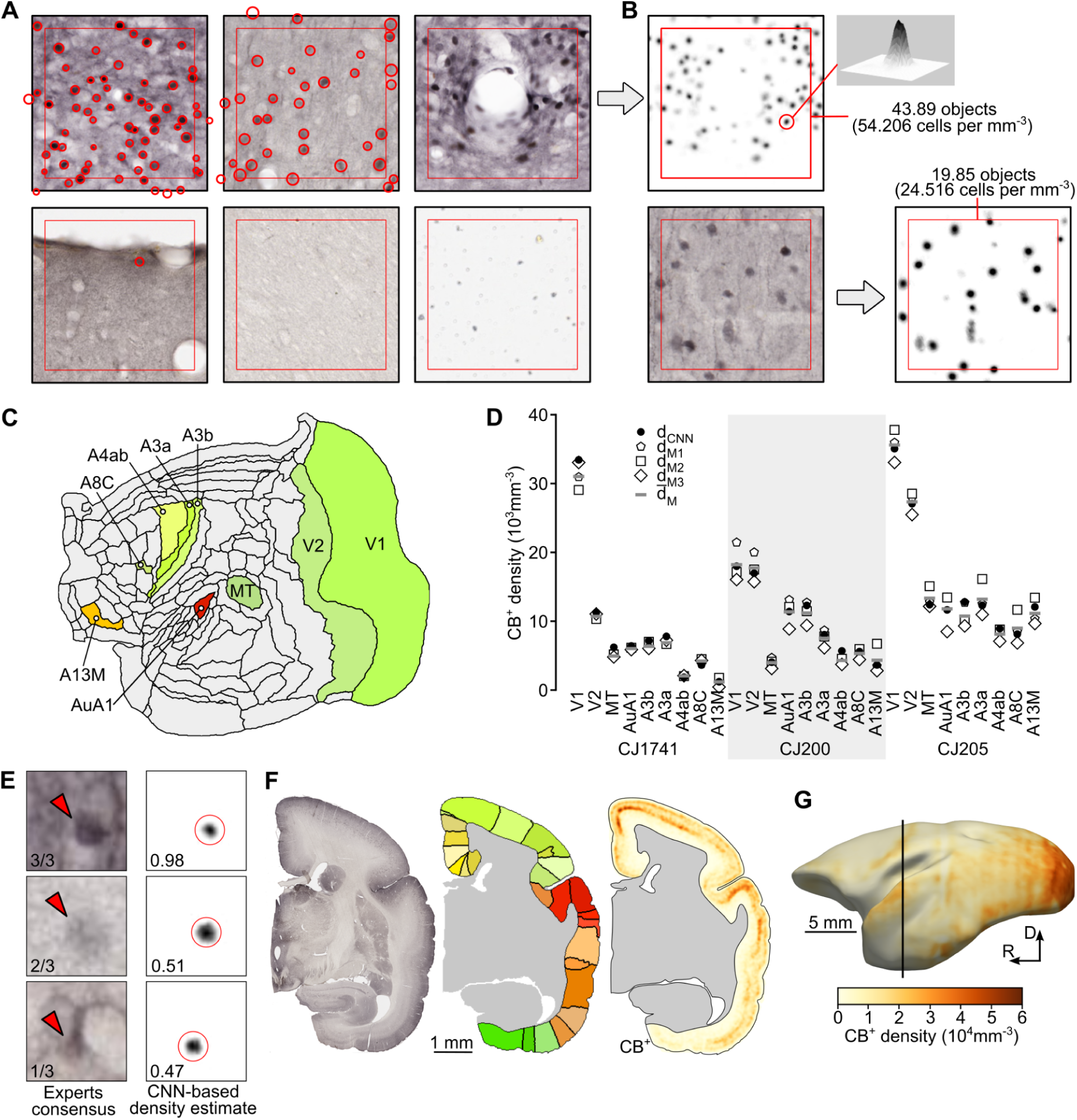
Training and evaluation of the U-Net Convolutional Neural Network (U-Net CNN). (A) Examples of counting boxes manually annotated by neuroanatomists (red circles of various sizes). The entire counting box is 150 × 150 µm while the area annotated with the red square is a 256 × 256 px (127 µm × 127 µm, 0.4974 µm per pixel) image patch used for training the CNN. A total of 4,072 counting boxes were defined, including boxes representing various densities of neurons *(top left* and *top middle)*, parts of the tissue containing blood vessels or artifacts *(top right* and *bottom left)*, as well as samples of white matter *(bottom center)* or samples that do not depict the brain tissue *(bottom right)*. (B) Density map generated based on the counting box indicated by the gray arrow. A single Gaussian blob corresponds to a single CB^+^ neuron, and 43.89 neurons are located within the indicated image patch (red square). Upon training, the U-Net CNN, a counting box not previously presented to the network, can be turned into a density map, and the total number of neurons within an image patch can be computed. (C) Comparison of the CB^+^ densities estimated by the U-Net CNN against multiple human annotators, illustrated for nine cortical areas (V1, V2, MT, AuA1, A3b, A3a, A4ab, A8C, and A13M, see Supplemental Table 1 for a list of areas and their abbreviations) that are unambiguously identifiable based on cyto- and myeloarchitecture, and which were sampled for benchmarking. (D) Per-area (i.e., average values for all boxes sampled from a given area) densities for the three analyzed hemispheres. Different symbols show the results for individual human experts (d_M1_ to d_M3_), an average of the three expert observers (*d̅*_M_), and the densities obtained with the U-Net CNN (d_CNN_). The mean of the differences between d_CNN_ and the d_M_ densities is statistically indifferent from zero (see Fig. S1F, G for statistical details), indicating that the automatic and the average manual counts are indistinguishable. (E) The results obtained by the U-Net CNN reflect the consensus between the expert annotators. (*rows, top to bottom*) Examples of individual CB^+^ neurons marked by all, two, and only one expert, respectively, and a proportional density estimate by the U-Net CNN. The proposed method helps alleviate the interindividual variability of manual cell counting. (F) We applied the procedures for estimating the density of CB^+^ neurons to all CB-stained sections in all three considered hemispheres. Here, results for an example section (CJ1741-r16c) are presented. From left to right: section image, segmentation of the cortex into individual areas based on manual parcellation and coregistration to the reference template (Majka et al. 2020, see *Methods* for details), and density map of CB^+^ neurons (see G for scale). Note that the quantification of the results is performed only in the cortical areas, while the subcortical regions are not considered. (H) Example three-dimensional reconstruction of a CB^+^ density map constituting the basis of the flatmap plots in the further figures. The black line indicates the location of section r16c presented in panel (F). The datasets are available for download from https://www.marmosetbrain.org/whole_brain_cb_maps.

### Areal distribution of CB^+^ neurons

Figures 3 and 4 show the density (neurons/mm^3^) of CB^+^ neurons across different cortical areas. For the summary in Fig. 3A, cortical areas were arranged in groups defined by location and function (Reser et al., 2013). The CB^+^ neuronal density varies according to area, within an approximately six-fold range (5·10^3^ and 30·10^3^ cells/mm^3^; means for 3 animals; individual data shown in Fig. 4). Given that the total neuronal density estimated by NeuN staining is also known to vary substantially between cortical areas (Atapour et al., 2019), a natural question is whether these observations can be explained by a simple model whereby CB^+^ neurons form a constant fraction of the neuronal population across areas. Our analyses demonstrate that this is not the case. Despite a significant correlation being found between the density of CB^+^ neurons and total neuronal density (Fig. 3B, top left), the data revealed that areas containing higher densities of CB^+^ neurons also tend to have these forming higher percentages of the total neuronal population (Fig. 3B, top right and bottom graphs show data from individual animals). Significant differences between areas in different functional groups persist after correction for total neuronal density (Fig. 3C). Thus, different cortical areas show variations in intrinsic cellular circuitry, suggesting different functional requirements mediated by CB^+^ neurons.

Auditory areas were particularly notable, as a group, for having high absolute and relative proportions of CB^+^ neurons, with the highest peak relative densities (CB^+^ neurons as a percentage of the total neuronal population) occurring in the auditory core areas (AuA1, AuR, AuRT). Visual areas also tended to show high absolute CB^+^ neuron densities (Figs. 3 and 4), but these are somewhat offset by high overall neuronal densities (Atapour et al., 2019), resulting in less notable relative densities (Fig. 3C). The primary visual cortex (V1) was an outlier both in terms of overall CB^+^ neuronal density, and in having these neurons forming a very high proportion of the total neuronal population (Figs. 3B, D; see Fig. 5 for individual data), with the second visual area (V2) providing a transition between the extreme values in V1 and those in other visual areas. Although the somatosensory cortex (SSC) included areas with a wide range of CB^+^ neuronal densities, the primary areas (areas 3a and 3b) represented well-defined local maxima, which contrasted with the adjacent motor/ premotor complex and posterior parietal areas (Figs. 3D and 5).

**Figure 3.**
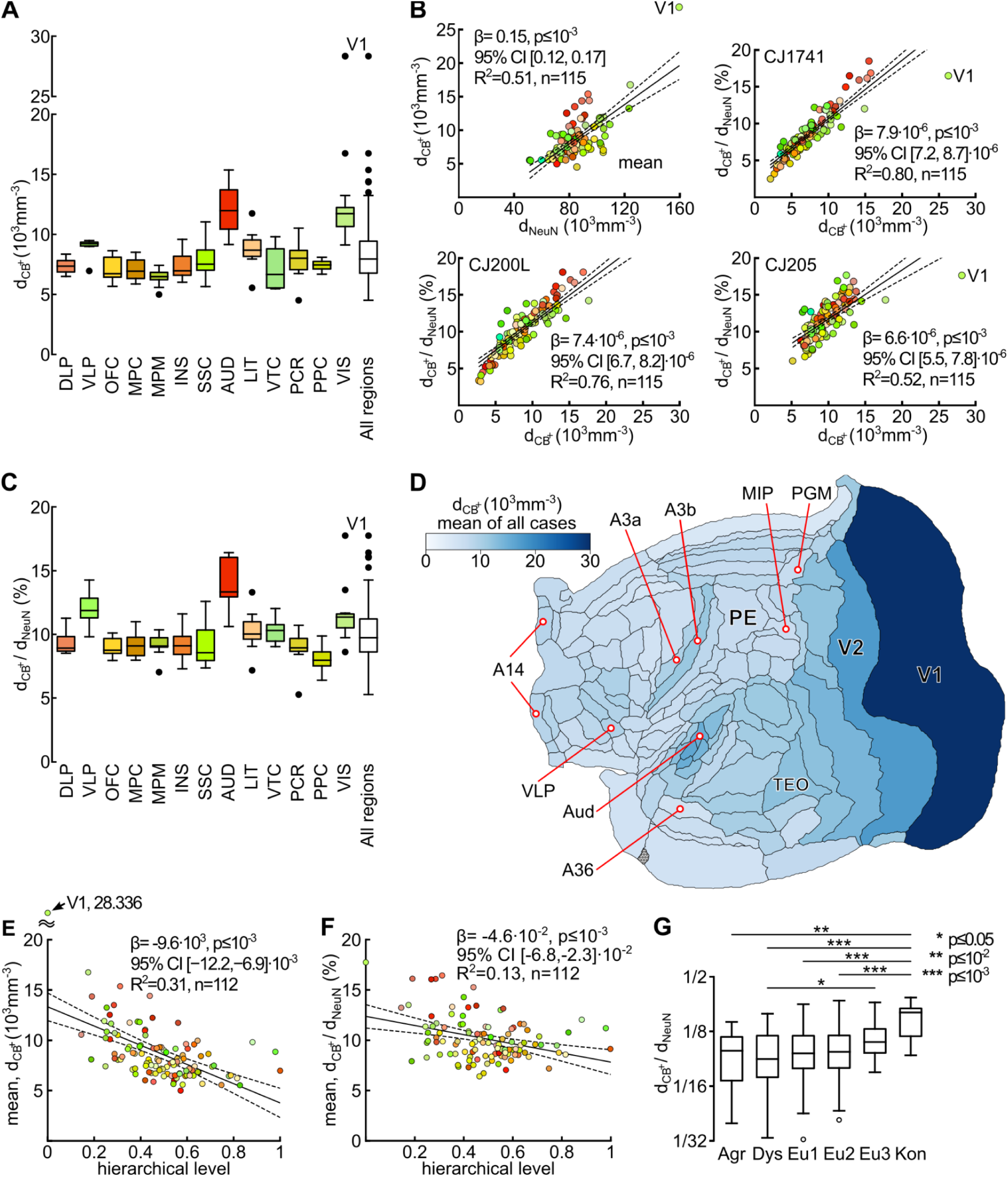
Variation in the density of calbindin-positive (CB^+^) neurons across the marmoset cortical areas. (A) Density (neurons/mm^3^) of CB^+^ neurons across areas, grouped according to the classification proposed by Reser et al. 2013. For each box plot, the center line indicates the median, the box limits the upper and lower quartiles, the whiskers 1.5× the interquartile range, and the annotated points outliers). Abbreviations: DLP: dorsolateral prefrontal cortex; VLP: ventrolateral prefrontal cortex; OFC: orbitofrontal cortex; MPC: medial prefrontal cortex; MPM: motor and premotor cortex; INS: insular cortex; SSC: somatosensory cortex; AUD: auditory cortex; LIT: lateral and inferior temporal cortex; VTC: ventral temporal cortex (encompassing parahippocampal, perirhinal and entorhinal areas); PCR: posterior cingulate and retrosplenial cortex; PPC: posterior parietal cortex; VIS: visual cortex. (B) Top left: the relation between the absolute density of CB^+^ neurons and total neuronal density (NeuN staining) in different areas. Top right and bottom: relations between relative and absolute densities of CB^+^ neurons in 3 animals. (C) Percentages of CB^+^ neurons across groups of areas, classified as in panel A. (D) Flat map representation of the CB^+^ neuronal density (mean of all 3 animals) in different areas. Some of the cortical areas are identified for orientation. (E, F) The relation between the absolute and relative densities of CB^+^ neurons and hierarchical level derived from laminar patterns of connections between cortical areas (Theodoni et al. 2021). (G) Differences in relative density of CB^+^ neurons between areas according to the degree of lamination (adapted from John et al. 2022). Abbreviations: Agr: agranular areas; Dys: dysgranular areas; Eu1, Eu2, Eu3: eulaminate areas with increasing levels of laminar differentiation; Kon: koniocortical areas. Colors in B, E, and F correspond to those in A and C.

**Figure 4.**
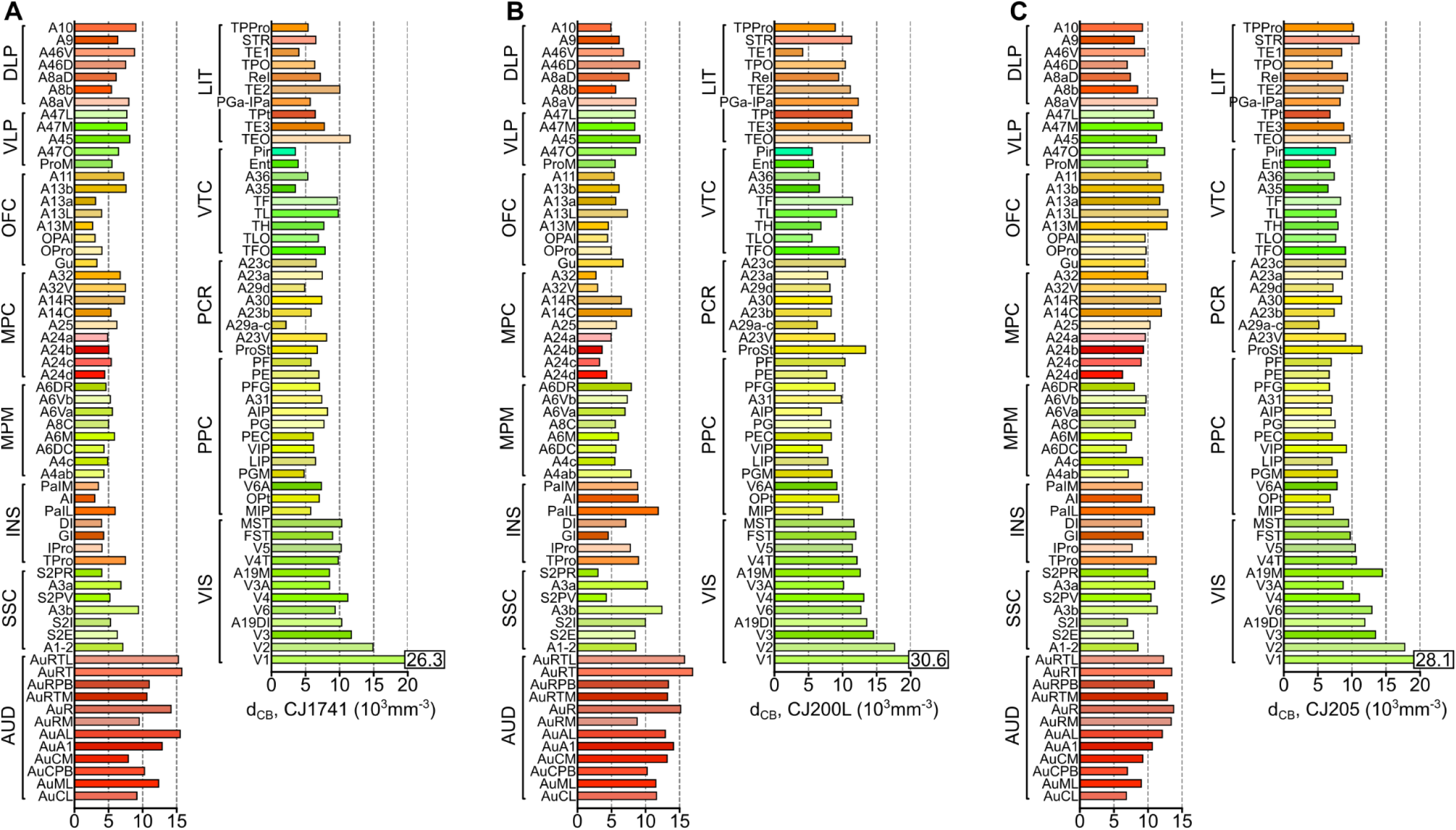
Densities of CB^+^ neurons in cortical areas of three marmosets. For this figure, areas were grouped according to the classification proposed by Reser et al. 2013 (modified from Paxinos et al. 2012). Abbreviations (groups of areas): DLP: dorsolateral prefrontal cortex; VLP: ventrolateral prefrontal cortex; OFC: orbitofrontal cortex; MPC: medial prefrontal cortex; MPM: motor and premotor cortex; INS: insular cortex; SSC: somatosensory cortex; AUD: auditory cortex; LIT: lateral and inferior temporal cortex; VTC: ventral temporal cortex (encompassing parahippocampal, perirhinal and entorhinal areas); PCR: posterior cingulate and retrosplenial cortex; PPC: posterior parietal cortex; VIS: visual cortex. The abbreviations of individual areas follow the nomenclature proposed by Paxinos et al. (2012).

**Figure 5.**
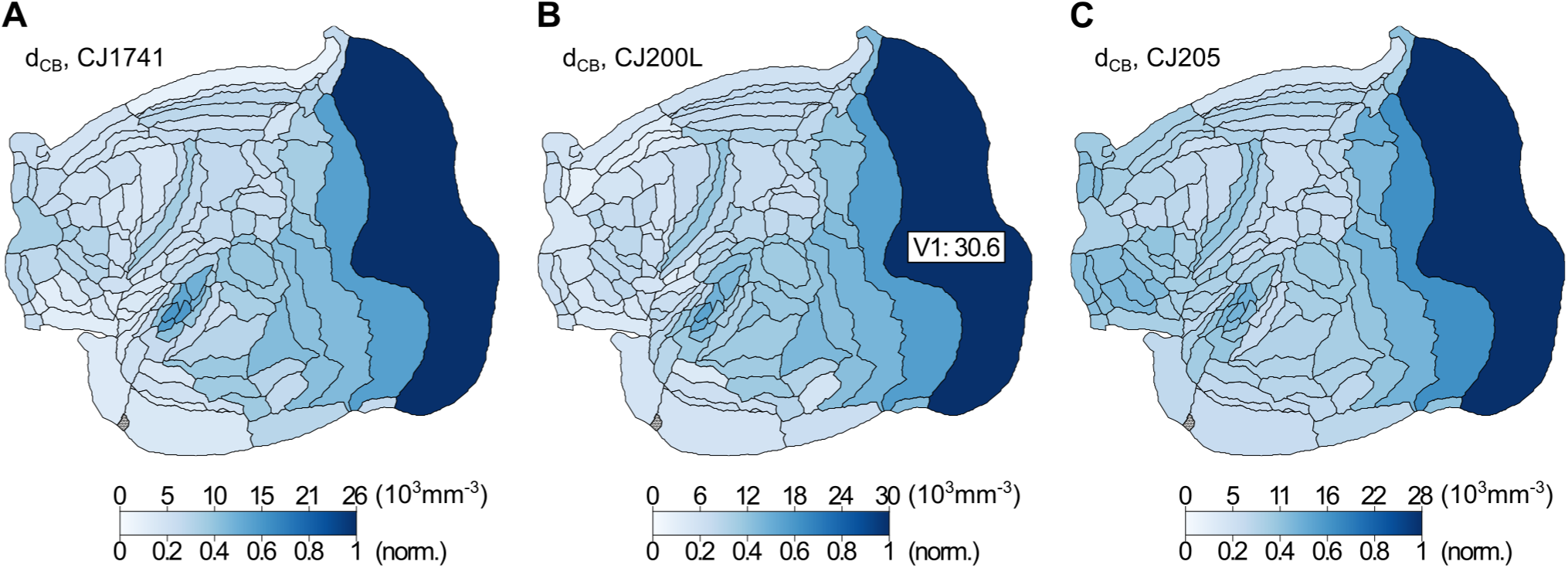
Maps of the distribution of CB+ neurons in three animals, shown in unfolded representations of a left hemisphere. For each map, the upper scale indicates densities in neurons/ mm3, and the lower scale shows the same data normalized to the peak density of a given individual.

Although isocortical association areas did not show extreme variations in CB^+^ neuronal density, a finer-grained analysis revealed a few notable trends. For example, in the prefrontal cortex, ventrolateral prefrontal areas (VLP) such as 45, 47M, and 47O tended to show higher percentages of CB^+^ neurons in comparison with other (frontopolar, dorsolateral prefrontal, orbitofrontal, and medial prefrontal) subdivisions (e.g. Fig. 3C). In the temporal lobe, caudal lateral and inferior temporal areas such as TEO, TE2, and TE3 showed higher absolute CB^+^ neuron densities in comparison with ventral (VTC) areas such as 36 and TH (Fig. 3A), but relative densities showed less variation (Fig. 3C). The posterior parietal areas (PPC) were distinct, as a group, for low relative densities of CB^+^ neurons (Fig. 3C).

### Correlation between the distribution of CB^+^ neurons and hierarchical levels

One of the basic organizational principles of the primate cortex is the existence of an anatomical hierarchy of areal connectivity, which can be quantified based on the laminar origins and terminations of cortico-cortical connections (Rockland and Pandya, 1979; Maunsell and Van Essen, 1983). This hierarchy, which is theorized to reflect the principal direction of information flow from sensory input to the generation of behavioral responses and executive function (Barone et al., 2000; Markov et al., 2014; Chaudhuri et al., 2015), has been determined for many areas of the marmoset cortex (Theodoni et al., 2021). This raises the question of whether there is a systematic relation between the density of CB^+^ neurons and the hierarchical rank. We found that hierarchically “low” areas, such as the primary sensory areas, tend to show high densities of CB^+^ neurons, compared to “high” areas (Fig. 3E). A similar but weaker relation is also detected when the percentages of CB^+^ neurons in different areas was considered (Fig. 3F).

Dombrowski et al. (2001) demonstrated that the distribution of cell types defined by expression of calcium-binding proteins varied according to the degree of differentiation between cortical layers (a characteristic that is hypothesized to reflect the evolutionary history of the mammalian cortex; Pandya et al., 2015) in areas of the macaque prefrontal cortex. When analyzed according to the six main cytoarchitectural categories of lamination proposed for the primate cortex (adapted for the marmoset cortex from the classification detailed by John et al., 2022), we found that CB^+^ neurons tended to form a higher proportion of the neuronal population in areas with the highest degree of lamination (eulaminate 3 and koniocortex), but few differences were apparent between other types of the cortex (Fig. 3G).

### Laminar distribution of CB^+^ neurons

Next, we addressed the extent to which the laminar distribution of CB^+^ neurons varied in different areas. To perform this analysis, the upper and lower boundaries of layer 4 were delineated in adjacent Nissl-stained sections, generating a mask that was transferred to the 3-dimensional reconstruction based on CB-stained sections. In a few isocortical areas where a clearly delineated layer 4 was not obvious (e.g. the representations of the limb and axial musculatures in the primary motor cortex, A4ab), we analyzed separately a thin (75 µm wide) strip of cortex centered on the interface between layers 3 and 5; areas without a defined layer 4 homolog (e.g. the entorhinal and piriform areas, see *Methods*) were not considered in this analysis. Other landmarks including the interfaces between layers 1 and 2 and between layer 6 and white matter were also incorporated, therefore allowing us to analyze the distribution of neurons according to three compartments in each area: layers 2 and 3 (supragranular), layer 4 (granular) and layers 5 and 6 (infragranular). To quantify areal differences we calculated the ratios of CB^+^ neuronal density in supragranular to infragranular layers (d_2,3_/ d_5,6_ ratio; Fig. 6A, C), and in supragranular to granular layers (d_2,3_/ d_4_ ratio; Fig. 6B, D).

**Figure 6.**
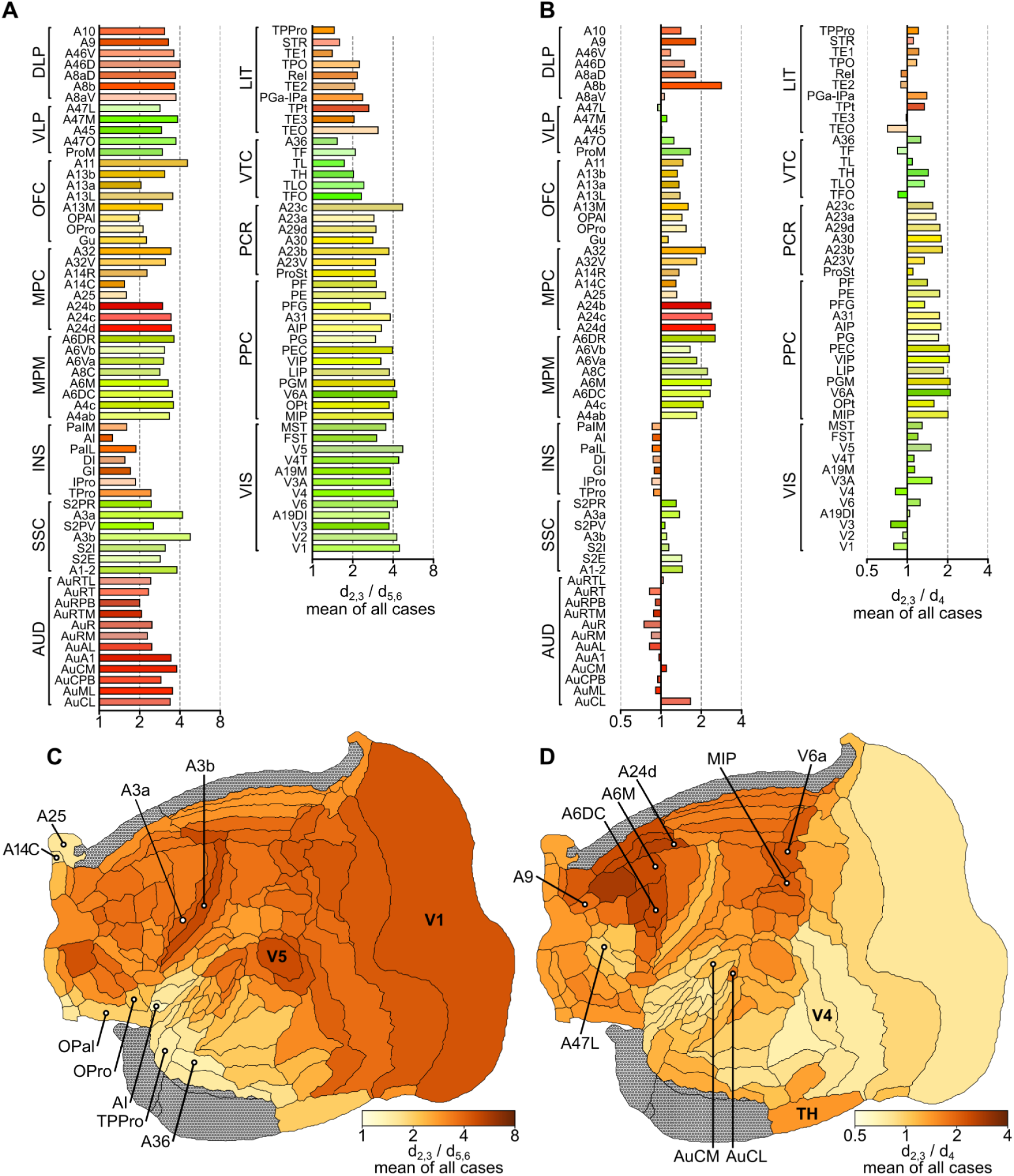
Laminar trends in the distribution of calbindin-positive (CB^+^) neurons in areas of the marmoset cortex. (A) Ratios of the density of CB^+^ neurons in supragranular versus infragranular layers (d_2,3_/ d_5,6_). (B) Ratios of the density of CB^+^ neurons in supragranular layers versus layer 4 (d_2,3_/ d_4_). (C, D) Flat map representations of the distributions of d_2,3_/ d_5,6_ and d_2,3_/ d_4_, summarizing the data shown in A and B. Areas without a clear homolog of layer 4 were excluded from analysis, indicated in gray.

Generally, CB^+^ neurons are more heavily concentrated in the supragranular layers throughout the cortex. As shown in Figure 6A, the ratio of CB^+^ neuronal density in supragranular to infragranular layers (d_2,3_/ d_5,6_ ratio) was higher than 1 in every area. However, the data also revealed significant differences between areas, with this ratio varying between 1.28 (agranular insula, AI) and 4.88 (area 3b). Lower d_2,3_/ d_5,6_ ratios (<2) were typically associated with areas of the limbic cortex, including most subdivisions of the insula (e.g. the agranular insula, AI), caudal orbitofrontal cortex (e.g. OPal, OPro), ventromedial frontal cortex (areas 14C and 25), the temporal pole (TPPro) and ventral temporal areas (e.g. area 36). In contrast, higher ratios (strong supragranular bias) were found in most visual areas and primary somatosensory areas (A3a and A3b), among others. Most of the areas of association isocortex and motor/ premotor areas (MPM) had intermediate d_2,3_/ d_5,6_ ratios, with CB^+^ neurons being 3-4 times as densely distributed in the supragranular layers, compared to infragranular layers. In addition, there was a strong negative correlation between CB^+^ neuron d_2,3_/ d_5,6_ ratio and hierarchical level (Fig. 7A), as well as a clear trend for an increase in this ratio with an increasing degree of lamination (Fig. 7B).

**Figure 7.**
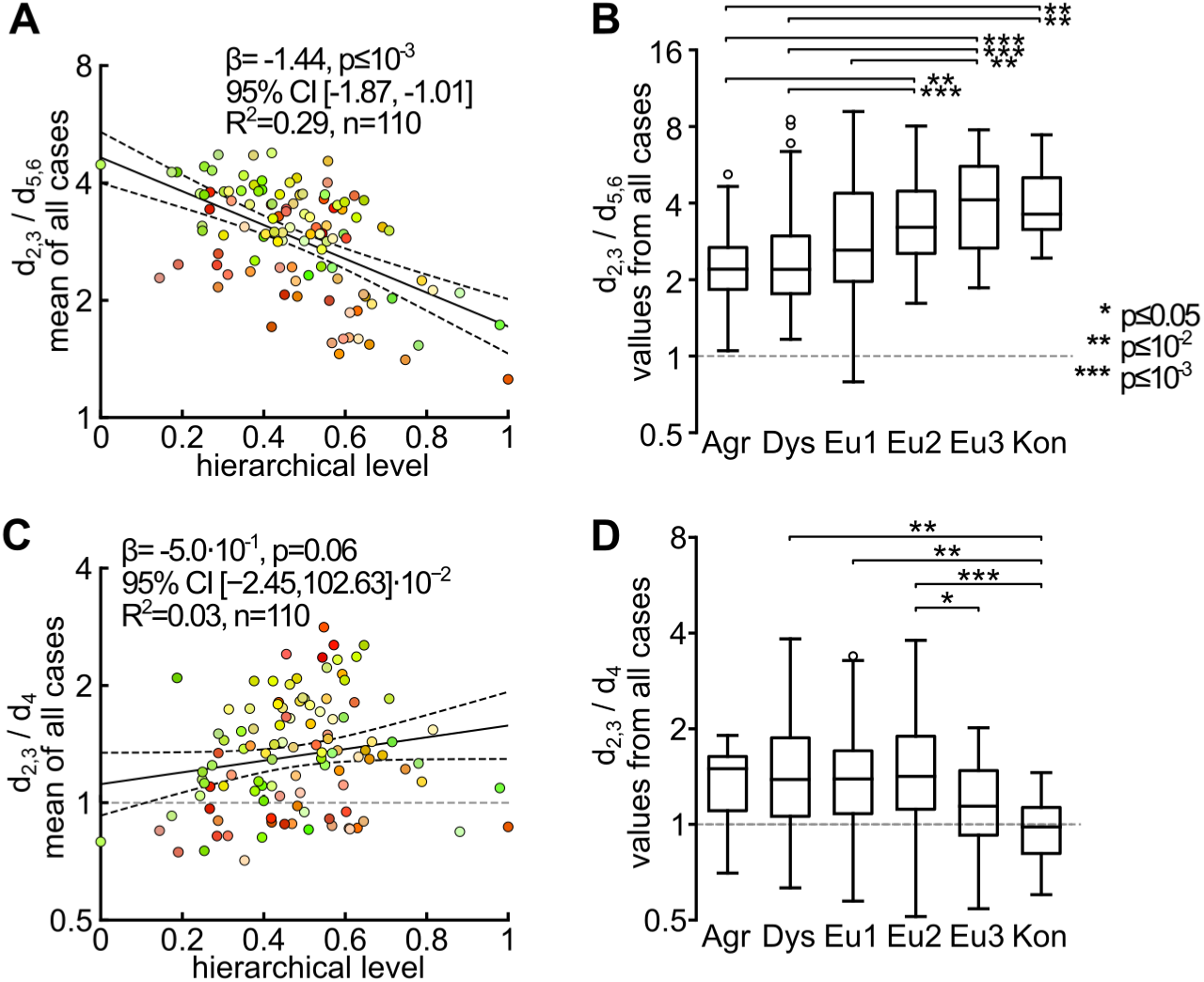
relations between the laminar distribution of CB^+^ neurons and large-scale trends in cortical organization. (A) The d_2,3_/ d_5,6_ ratios were strongly correlated with the area’s hierarchical level, derived from the laminar pattern of connections. This demonstrates that the bias towards higher density of CB^+^ neurons in supragranular layers was most marked in low-level (e.g. purely sensory) areas, and more subtle in association and premotor areas. (B) In parallel, the d_2,3_/ d_5,6_ ratios varied systematically with respect to the classification of an area according to the degree of laminar differentiation. Analyses of the laminar distribution of CB+ neurons. (C) In contrast with (A), there was no significant correlation between d_2,3_/ d_L4_ and hierarchical level. (D) Analysis of d_2,3_/ d_4_ relative to the degree of cortical lamination indicated significantly lower d_2,3_/ d_4_ ratios (i.e. higher relative concentrations in layer 4) in koniocortex and eulaminate 3 areas, compared to others.

There were, in addition, systematic differences between areas in terms of the ratio of CB^+^ neurons densities in layers 2-3 and layer 4 (d_2,3_/ d_4_). Here, the data revealed that in visual, auditory, and somatosensory areas the density of CB^+^ neurons in layer 4 tended to be similar to that found in the supragranular layers (Fig. 6B) resulting in relatively low d_2,3_/ d_4_ ratios. In contrast, CB^+^ neurons tended to be sparsely distributed around layer 4 in premotor areas (e.g. A6M, A6DC), cingulate motor areas (e.g. A24d) and posterior parietal areas that have known functions in visuomotor and somatomotor planning (e.g. MIP, V6a), resulting in much higher d_2,3_/ d_4_ ratios. This functional association was also suggested when visual and auditory areas belonging to different functional streams were considered. For example, visual areas linked primarily to the ventral stream (e.g. V4, TEO, TE3) all had d_2,3_/ d_4_ ratios <1, thus resembling V1 and V2 in having relatively high densities of CB^+^ neurons in layer 4 (Fig. 6B). In contrast, dorsal stream areas (e.g. V3A, V5, V6, MST, and FST), which provide critical input for the frontoparietal network (and thus for visual control of action; Goodale and Milner 1992; D’Souza et al., 2021), were all characterized by d_2,3_/ d_4_ ratios >1. Following a similar trend, whereas most of the auditory cortex areas had d_2,3_/ d_4_ ratios below 1, the exceptions were caudal auditory belt areas (AuCL, AuCM), which are part of the auditory dorsal (“where”) stream (Rauschecker et al., 2000). An analogous functional distinction in CB^+^ neuron distribution extended even to the prefrontal cortex, where ventrolateral areas (e.g. subdivisions of area 47), theorized to represent extensions of the visual and auditory ventral streams (Poremba and Mishkin, 2007; Diehl and Romanski, 2014), all showed low d_L23_/ d_L4_ ratios in comparison with dorsolateral areas associated with the dorsal streams (e.g. areas 8b, 8aD, 9 and 46; Kravitz et al., 2011).

The correlation of d_2,3_/ d_4_ with an area’s level in the anatomical hierarchy was not significant (Fig. 7C). Likewise, an analysis relative to the degree of cortical lamination indicated only a few differences, with the exception of lower d_2,3_/ d_4_ ratios (i.e. higher relative concentrations in layer 4) in koniocortex (Fig. 7D). In summary, the relative paucity of CB^+^ neurons in layer 4 appears to be more correlated with an area’s function (in particular its involvement in sensorimotor integration, or planning of action) rather than its relation to structural attributes of the cortex.

## Discussion

The incorporation of neuroinformatics to the discipline of neuroanatomy has been most evident in the rapid progress in our knowledge of the human brain, including the combination of results from non-invasive techniques which have allowed new insights at millimeter to centimeter scales (e.g. Elam et al., 2021). More recently, comparable change has been taking place in our knowledge at the level of cells and their connectivity, primarily using animal models. Starting with studies in rodents (Oh et al., 2014; Kim et al., 2017; Bjerke et al., 2020), the traditional approach of investigating areas and nuclei one or a few at a time is being complemented by projects involving larger datasets, and focused on unveiling organizational principles, often relying on multiple techniques. Such approaches have recently made their way to the brains of non-human primates, revealing new information based on single-neuron resolution connectivity (Majka et al., 2020; Skibbe et al., 2023) and transcriptomics (Krienen et al., 2020, 2023; Chen et al., 2023), sometimes combined with non-invasive imaging (Liu et al. 2020; Tian et al., 2022). However, one missing link in these efforts has been their relation to the traditionally recognized cell types of the mammalian cortex, for which a wealth of cytological and physiological knowledge is available, and which are central to the generation of biophysically realistic models of brain function (e.g. Wang et al., 2020). The present paper demonstrates a strategy to overcome this gap. Using a combination of traditional immunohistochemistry, expert annotation, validation by stereology, and neuroinformatics solutions (the latter aimed at high-throughput image analysis, brain reconstruction and cytoarchitecture-aware registration), we created the distribution maps of neurons defined by preferential expression of a calcium-binding protein across the entire marmoset cortex.

The *density* or *regression counting* paradigm employed in this study allows for estimating the total number of labeled cells within a defined region (e.g. a counting box) instead of identifying individual instances of CB^+^ neurons. This approach significantly reduces the annotation time, which is beneficial when objects are densely packed, occlude to a high degree, a large portion of tissue is to be analyzed, or the imaging data has a resolution insufficient to visualize the neuronal morphologies in an adequate level of detail. From this perspective, the proposed approach is affordable in terms of ease of implementation, modest annotation workload, and fast execution in production. Therefore, it can be more easily applied to the brain in less commonly used species without involving genetic modifications (Kim et al., 2017) or large-scale spatial transcriptomics (Chen et al., 2023). Another advantage is that, by relying on the strong expression of a protein, it can be more directly related to traditionally recognized cell types, about which there is a wealth of physiological information that can be leveraged for biophysical modeling (Ascoli et al., 2008; Markram et al., 2015; Tripathy et al., 2015).

Expert annotators exhibit a natural bias (e.g. Jacobs et al., 2001; Bloom and Harrington, 2004) and their annotations vary between each other or over time (i.e. between plotting sessions). Our approach, however, represents a consensus among human annotators in the dataset used in this study (as shown in Figure 2D and E), thereby helping to mitigate these biases. Evaluation of the model’s performance showed that the discrepancy between the manual and U-Net CNN counts reduces with the amount of tissue under consideration (Fig. S1F, G), see *STAR* ★ *METHODS, Comparison against multiple human raters*.

Alternative approaches involve object detectors based on classical machine learning methods, such as random forests (e.g. Berg et al., 2019; Bouvier et al., 2021) or deep learning instance segmentation architectures like those available in Detectron2 library (https://github.com/facebookresearch/Detectron) or Cellpose package (Stringer et al., 2021, https://www.cellpose.org/). The latter, however, require extensive and well-curated outlines of individual cells, which could potentially offset the cost reduction promised by the machine learning approaches in the first place. To encourage the development of improved methods for identifying and exploring CB^+^ density patterns, we have released the dataset used to train our model as a supplemental material. This dataset includes individual counting strips and annotations into boxes and marked cells (Fig. 1B-E).

Finally, we provide the results not only in tabular, but also in three-dimensional form. This enables wider interoperability, for instance warping the CB^+^ neuronal density maps into the space of MRI marmoset brain templates (e.g. Liu et al., 2020), and analyzing the density patterns in the context of structural and functional connectivity.

The present method has, therefore, the capacity to obtain cellular distribution maps for the whole cortex of single individuals, hence robust detection of quantitative differences between areas. This is significant given that there can be method-related differences in estimates of the prevalence of the same cell type obtained by different laboratories, in different individuals (Bjerke et al. 2020). Moreover, although the estimates of absolute cell densities can vary between individuals, the relative density demonstrates a highly reproducible pattern (Fig. 5). It remains unclear whether individual variation in the absolute density of CB^+^ neurons reflect methodological factors, or genuine (biologically relevant) differences, such as those related to sex (Abel et al. 2011), age (Bu et al. 2003) or postnatal experience. This is an important question, which will require systematic analysis of a much larger number of individuals.

The only previous study which has attempted an extensive mapping of CB^+^ neurons in the cortex was that of Bjerke et al. (2020), who studied the mouse brain. The distribution reported in this species is significantly different from the present results obtained in a non-human primate. For example, a significant peak in CB^+^ neuronal density was observed in the infralimbic area, which was not apparent in the corresponding region of the marmoset brain (subdivisions of medial prefrontal cortex). In addition, particularly low densities were observed in retrosplenial and posterior cingulate areas, which were unremarkable in this respect in the marmoset brain. Furthermore, the densities in auditory, visual and somatosensory areas were similar in the mouse, in contrast with differences observed in the marmoset. However, in both species relatively low densities of CB^+^ neurons were observed in motor and premotor areas. Finally, the absolute CB^+^ neuron densities were much lower in the mouse cortex (∼300 - 3500 cells/ mm^3^) compared to marmoset (∼5000 - 25000 cells/ mm^3^) or estimates obtained in the macaque frontal lobe (Dombrowski et al. 2001).

Although there have been previous reports of diversity in CB^+^ neuron distribution across areas of the primate cortex (e.g. Condé et al. 1994; Dombrowski et al. 2001; Elston et al. 2003; Bourne et al. 2007), the present results establish that these differences are not only of a larger magnitude, but also systematic in ways not previously appreciated. For example, auditory areas are notable in their high densities of CB^+^ neurons, and there were cortex-wide correlations between the prevalence of these cells and both hierarchical levels, defined by laminar pattern of connections, and type of lamination. These observations, which do not simply reflect differences in overall neuronal density, are compatible with the notion that sensory cortex (in particular, primary sensory areas) differs from association and motor planning areas in terms of intrinsic circuitry, including the required levels of GABAergic inhibition.

This heterogeneity is also underlined by systematic differences in laminar distribution. Although CB^+^ neurons are overall more densely distributed in the supragranular and granular layers, this trend was more pronounced in sensory areas, and least pronounced in limbic areas. In addition, across the entire cortex areas involved in sensorimotor integration and action planning showed reduced densities of CB^+^ neurons in layer 4, compared to areas more directly related to sensory analysis. This correlation is independent of the hierarchical levels at which such function is performed, or the presence of a clearly defined layer 4 (e.g., it is also evident in posterior parietal areas). These observations may reflect the temporal level of integration needed for different functions (Li and Wang 2022), with the shorter time scale required for visual and auditory analysis requiring more precise spatiotemporal modulation of the levels of inhibition. Together, the present results significantly extend our knowledge of the degree of heterogeneity across areas of the primate cortex, which will need to be considered by future models of cortical function.

## Conclusions

CB^+^ neurons have long been recognized as one of the main categories of neurons in the mammalian cortex, but little has been known about the principles governing their distribution relative to different functional systems, or, more generally, in the primate brain. Enabled by computational techniques that allow whole-cortex mapping, the present results highlight a previously unsuspected degree of heterogeneity across cortical areas and layers, and significant differences relative to data obtained in rodent brains. These observations further highlight the need for work aimed at establishing a firm basis for accurate biophysical models of cortical function, and hence meaningful attempts to simulate neural activity related to different sensory, motor or cognitive functions.

## Limitations of the study

The primary limitation of the present approach is that it cannot differentiate between neurons that share preferential expression of a protein, but may be physiologically distinct. The focus of the present study was to establish density maps of CB^+^ neurons rather than analyze their subtypes. However, this limitation can be at least partially addressed in future work by using deep learning architectures capable of instance segmentation (e.g. Kayasandik et al., 2020; Wu et al., 2022) to distinguish morphological categories (e.g. Xiong et al., 2022). Additionally, adopting higher imaging magnifications (e.g. García-Cabezas et al., 2016) combined with double-labeling for other molecular, markers and followed by generating expert-curated training sets, could provide further insights on the distribution of more specifically defined cell categories.

## Author contributions

Nafiseh Atapour: Data Curation, Formal Analysis, Methodology, Validation, Writing – Original Draft Preparation, Writing – Review & Editing.

Marcello G.P. Rosa: Conceptualization, Funding Acquisition, Methodology, Project Administration, Validation, Supervision, Writing – Original Draft Preparation, Writing – Review & Editing.

Shi Bai: Data Curation, Software, Writing – Review & Editing

Sylwia Bednarek: Formal Analysis, Software, Writing – Review & Editing. Agata Kulesza: Formal Analysis, Visualization.

Gabriela Saworksa: Formal Analysis, Visualization.

Katrina H Worthy: Methodology, Project Administration

Piotr Majka: Conceptualization, Data Curation, Formal Analysis, Funding Acquisition, Methodology, Project Administration, Software Supervision, Validation, Visualization, Writing – Original Draft Preparation, Writing – Review & Editing.

## Acknowledgements

Funding was provided by the National Science Centre (2019/35/D/NZ4/03031 to Piotr Majka), the National Health and Medical Research Council to Marcello G. P. Rosa (APP1194206) and Nafiseh Atapour (APP2019011).

The authors would like to acknowledge the contributions of Cecilia Cranfield, Daria Malamanova and Melissa Chong during the data input phase (manual annotation of images). We also thank Emilia Chojak and Piotr Szulim for assistance in data quality assurance and image processing.

## Declaration of interests

The authors declare no competing interests.

## STAR ★ METHODS

### KEY RESOURCES TABLE

| REAGENT or RESOURCE | SOURCE | IDENTIFIER |
| --- | --- | --- |
| **Antibodies** |  |  |
| Mouse monoclonal (IgG1) anti-Calbindin-D28K | Swant Inc, Swiss | Code No. 300<br>RRID: AB_10000347 |
| **Biological samples** |  |  |
| Three healthy adult marmoset brains | Monash Animal Research platform (MARP) | https://research.monash.edu/en/equipments/animal-research-platform-marp |
| **Critical commercial assays** |  |  |
| Biotinylated horse anti-mouse IgG secondary antibody | Vector Laboratories | PK-6102<br>RRID: AB_2336821 |
| DAB substrate | Vector Laboratories | SK-4100<br>RRID: AB_2336382 |
| **Deposited data** |  |  |
| Dataset used for training and validation of the Unet CNN | This paper | https://www.marmosetbrain.org/whole_brain_cb_maps |
| CB+ density maps, segmentations into cortical areas and layers | This paper | https://www.marmosetbrain.org/whole_brain_cb_maps |
| Nencki-Monash marmoset brain template | https://www.marmosetbrain.org/nencki_monash_template | RRID:SCR_018367<br>https://doi.org/10.1016/j.neuroimage.2020.117625 |
| **Experimental models: Organisms/strains** |  |  |
| *Callithrix jacchus* | Monash Animal Research platform (MARP) | https://research.monash.edu/en/equipments/animal-research-platform-marp |
| **Software and algorithms** |  |  |
| Custom Python scripts for training the Unet CNN | This paper | Supplemental materials,<br>https://www.marmosetbrain.org/whole_brain_cb_maps |
| Advanced Normalization Tools (ANTs), version 2.1-linux | http://stnava.github.io/ANTs/ | RRID:SCR_004757<br>https://doi.org/10.1016/j.neuroimage.2014.05.044 |
| ITK-SNAP, version 3.8.0 | http://itksnap.org | RRID:SCR_00201<br>https://doi.org/10.1016/j.neuroimage.2006.01.015 |
| Aperio ImageScope | Leica Biosystems | https://www.leicabiosystems.com/en-au/digital-pathology/manage/aperio-imagescope/ |

### RESOURCE AVAILABILITY

#### Lead contact

Further information and requests for resources and reagents should be directed to and will be fulfilled by the lead contact, Piotr Majka.

#### Materials availability

This study has generated new materials in the form of datasets listed under the *Deposited data* section of the Key Resources Table.

#### Data and code availability

1. The density maps and the segmentations into cortical areas and layers for each case are available as Supplemental NIFTI files (available for download from https://www.marmosetbrain.org/whole_brain_cb_maps).
2. Training datasets in the form of annotated counting strips, as presented in Figure 1, are available as TIFF files containing the imaging data and corresponding SVG files containing the annotations. The dataset can be downloaded from https://www.marmosetbrain.org/whole_brain_cb_maps.
3. Summarized results that allow one to reproduce the analyses presented in this article are provided in Supplemental Table 1.
4. Custom Python scripts for training the Unet CNN are available as Supplemental Materials.
5. Any additional information is available from the lead contact upon request.

### EXPERIMENTAL MODEL AND SUBJECT DETAILS

This project was reviewed, approved, and monitored by the Monash University Animal Ethics Committee (Project ID: 26071;” Enabling an accurate model of brain wide connections”. After experiments unrelated to the present study, three marmosets (*Callithrix jacchus*, aged between 41–43 months, including two females [CJ1741 and CJ200L] and one male [CJ205]) were overdosed using sodium pentobarbitone (100 mg/kg, i.m.), and perfused transcardially using heparinized saline, followed by 4% paraformaldehyde in 0.1 M phosphate buffer (PB). The collected brains were post-fixed and cryoprotected in a buffered paraformaldehyde solution containing increasing concentrations of sucrose (10%, 20%, and 30%). The intact hemisphere was sectioned at 40 µm, yielding five sequential series of coronal sections used for different stainings. Myelin and Nissl-stained series of sections were used to delineate area boundaries (for details, see Atapour et al., 2019; Majka et al., 2020), with the Nissl-stained sections being in addition used to define the cortical layers (see below). Another series was used for CB immunostaining, as described previously in detail (Chong et al., 2022). Briefly, the sections were incubated in blocking solution (10% normal horse serum and 0.3% Triton-X100 in 0.1 M PB) for one hour at room temperature before undergoing primary antibody (calbindin-D28K, 1:8000, Swant Swiss, Code No. 300, RRID: AB_10000347) incubation at 4°C. A biotinylated horse anti-mouse IgG secondary antibody (1:200, PK-6102, Vectastain elite ABC HRP kit, Vector Laboratories, RRID: AB_2336821) incubation was then conducted for 30 min - followed by treatment with Avidin-Biotin Complex (ABC) reagent (PK-6102) and DAB substrate working solution (DAB kit SK-4100, RRID: AB_2336382) as a chromogen. Stained sections were scanned using an Aperio Scanscope AT Turbo system (Leica Biosystems), providing a resolution of 0.4974 μm per pixel.

### METHOD DETAILS

#### Manual annotation of CB^+^ neuronal somata

The manual cell annotation was performed using an in-house web-browser-based interface which enabled a human annotator to flexibly browse the microscopic resolution images, place counting boxes, and identify individual neuronal somata. Neuroanatomists could then manually annotate the location of every neuron within a counting box by placing a circular marker centered on the soma, and then adjusting the marker radius to reflect the size of the neuronal body (Fig. 2A).

In case CJ1741, following reconstruction and registration (see *Three-dimensional reconstruction,* below), columns of counting boxes, each 150 µm × 150 µm, were defined for each of the 116 areas, to encompass all cortical layers and a fragment of underlying white matter (Fig. 1B, C). Further, additional boxes were placed liberally to increase the diversity of the image features (e.g. blood vessels, artifacts, see Fig. 1D, E) to improve the training of the network. This dataset, comprising 2,220 counting boxes was split randomly into training (1,617 counting boxes) and validation (603 counting boxes) batches.

The training dataset included boxes from at least one strip for each of the 116 areas from case CJ1741 (Fig. 1B). These were further complemented with an auxiliary dataset comprising boxes covering the subcortical white matter and parts of the images that did not depict brain tissue (1,812 boxes, in total). These boxes were inspected visually to ensure they contained no CB^+^ neurons (Fig. 2A, bottom middle and bottom right). Subsequently, they were included in training dataset to suppress false positive rate by presenting the neural network examples of cell-alike objects such as dust and spots, smudges, etc.

In addition, in all three examined hemispheres, in nine selected areas representing various types of cytoarchitecture (V1, V2, MT, AuA1, A3b, A3a, A4ab, A8C, and A13M; Fig. 2C), additional strips were defined and annotated independently by three experts. This dataset was used to assess the performance of the automated method against multiple human raters (see *Comparison against multiple human raters,* below).

#### Automated detection of CB^+^ neurons

The automated estimation of densities of CB^+^ neurons relied on the *regression* or *density counting* approach (Sindagi et al., 2018; Xie et al., 2018; Hoekendijk et al., 2021). This paradigm allows for estimating the *total number* of labeled cells within a defined region (e.g. counting box) of the microscopic images, in contrast to identifying individual instances of cells. Since the *regression counting* has a statistical nature, the number of detected cells might be expressed as a non-integer value. Our solution uses the U-Net (Ronneberger et al., 2015) convolutional neural network (U-Net CNN), derived from Xie et al., (2018) implementation, to map an input microscopic, color (24 bits per pixel, RGB) image of an immunohistochemically stained tissue into a density map of CB^+^ neurons (Fig. 2B).

For each 150 µm × 150 µm (302 px × 302 px) counting box, a corresponding ground truth density map image was generated based on markers placed by the human annotators (Fig. 2B). Each marker was converted into a 2D Gaussian blob of a size proportional to the radius of the marker. This way, the entire blob sums up to 1.0 (a single neuron); hence the sum of all blobs in a counting box corresponds to the total number of neurons identified within this box. To avoid edge effects and to increase the training performance, the counting box image and the density map were cropped to 256 px × 256 px (127 µm × 127 µm), hereafter referred to as *image patches* (Fig. 2A, B).

In summary, the training dataset comprised 3,429 pairs of image patches and corresponding density maps, of which 64.5% (2,216) contained no CB^+^ cell annotation. In the remaining 1,213 boxes, the median number of neurons was 4.9, while the total was 9,219. The validation dataset consisted of 603 pairs of image patches and density maps totaling 3,072 neurons (a median of 4.78 neurons per patch), while 109 density maps contained no CB^+^ cell annotation.

#### Training of the U-Net CNN

Each of the red, green, and blue channels of an image patch was independently normalized to zero mean and a unit standard deviation. The normalized image patches and corresponding density maps were then piped into four-level-deep U-Net CNN. On each level, the convolutional layers were set to 64 filters, 3 px kernel size, and ReLu activation, while the final convolutional layer was set to linear activation. The model was trained for 160 epochs with varying learning rates: 10^-2^ (iterations 1–40), 10^-3^ (iterations 41–80), 10^-4^ (iterations 81–120), and 10^-5^ (iterations 121–160). Image augmentation included intensity and contrast variations, 0–45° rotation, and random horizontal and vertical flips. As the purpose of the network is to map (regress) an image into a density map, the mean square error (MSE) was used as the loss function, and the mean absolute error (MAE) was calculated for monitoring.

The U-Net CNN returns a total number of CB^+^ neurons within a given image patch (Fig. 2B). To calculate the density (i.e., number of objects per unit of physical volume), the following conversion is applied: d = 0.801·n·s^-2^·t^-1^, where *d* is the density, *n* is the number of neurons detected within an image patch, *s^2^* is the surface of a counting box (127 µm × 127 µm), *t* is the nominal thickness of the section (40 µm), and the factor of 0.801 is applied to correct for the shrinkage (Atapour et al., 2019).

#### Training results

The network successfully learned to map the images into CB^+^ density maps. A strong linear relation between the ground truth and U-Net CNN-based counts (n_m_= 1.021·n_CNN_ + 0.299, R^2^=0.896) can be observed (Fig. S1A), and the distribution of the residuals averages to zero. The relative difference between the automated count and the baseline decreases with the density (Fig. S1B). The CNN learned well to avoid spurious objects as the median error for a validation box that is known to contain no cells is only 0.06 (i.e. given an empty image patch, the U-Net CNN estimate is going to be <70 cells per mm^-3^ for the half of the patches, Fig. S1C).

#### Comparison against multiple human raters

Since the model was trained on samples derived from a single marmoset hemisphere (CJ1741) manually annotated by a single expert neuroanatomist, there could be a risk of overfitting the model, which could negatively affect the performance on the hemispheres that did not contribute to the training process. To assess the model’s accuracy on samples from the other cases (CJ200L and CJ205) and to confront the U-Net CNN performance against multiple experts, we generated a holdout (benchmark) dataset. In all three analyzed hemispheres, nine cortical areas (V1, V2, MT, AuA1, A3b, A3a, A4ab, A8C, and A13M; Fig. 2C) were selected. In each area, a single strip was defined and then annotated independently by three neuroanatomists. Therefore, the benchmark dataset amounts to 313 counting boxes per expert (939 in total). The counting boxes were pre-processed and converted into benchmark density maps as described above (Fig. 2B) and were processed by the U-Net CNN. Finally, the neuronal densities were calculated based on both sets of density maps: computed with the U-Net CNN and those defined by the experts.

A clear linear relation between the ground truth and CNN-based counts was found (d_M_=1.093·d_CNN_ - 0.90, R^2^=0.938, Fig. S1D). The U-Net CNN results seem slightly but systematically lower than the manual estimates (95% CI: [1.07–1.11]). Analysis of the discrepancy against the neuronal density (Fig. S1E) reveals that the primary sources of the differences are the counting boxes with the highest densities (>70·10^3^ cells per mm^-3^, see also the red rectangle in Fig. S1D), found in layer 4 of areas V1 and V2. For these densities, the automated results can be noticeably lower; for the remainder of the boxes, the automated and manual results match well.

The results also show that the densities obtained by human experts carry a noticeable variability (e.g. Fig. 2D). Since we consider each expert’s results equally important, we averaged (*d̅*_M_) the densities obtained by each human annotator (d_M1_ to d_M3_) for each counting box, and analyzed those densities against the U-Net CNN. First, we found that the mean U-Net CNN density for each area (i.e. the average density of all counting boxes in a given cortical area) and its manual counterpart are statistically indifferent in either absolute or relative terms (Fig. S1F, G white violin plots). This is also the case when it comes to the individual counting boxes. However, the spread of the differences is much more prominent (Fig. S1F, G violin plots in gray).

In the case of per-area estimates, the distribution of differences between the manual and the U-Net CNN densities is normal in either absolute or normalized terms (Shapiro–Wilk test: p=0.22 and p=0.79, respectively), and the average difference is indifferent from zero (t-test p values of 0.13, and 0.11 for absolute and normalized differences, respectively). As far as image patch (127 µm × 127 µm) densities are concerned, the distribution of both the absolute and normalized differences is not normal (Shapiro–Wilk test: p<10^-5^ in both cases), yet the differences are statistically indifferent from zero (Wilcoxon signed-rank test p=0.12 and p=0.07, respectively). In other words, the variance of the differences is noticeable, yet, on average, the U-Net CNN and manual results are comparable. When several counting boxes are considered (e.g. a set of boxes that belong to a single cortical area), the variance of the differences is noticeably smaller (Levene test p-values of 0.016 and 6·10^-4^ for differences of variances between the differences of areas and boxes, absolute and normalized, respectively), with the differences still statistically indifferent from zero. In conclusion, the U-Net CNN estimates of CB^+^ neurons reflect the average of human experts well, particularly when an area equivalent to several counting boxes is considered (∼0.25 mm^2^, or more).

#### Three-dimensional reconstruction

To segment the individual hemispheres into cortical areas, we performed a computational alignment and three-dimensional reconstruction following procedure detailed in Figures 4 and 5 in Majka et al., (2016) and Supporting Figures 5 and 6 in Majka et al., (2020). As a reference, we used the Nencki-Monash marmoset brain template (Majka et al., 2021; RRID:SCR_018367), which represents an average morphology of twenty young adult individuals of similar age range as the individuals used in this study. This helps, to a large degree, to mitigate the issue of interindividual variability. Since the Calbindin-stained sections were used instead of Nissl-stained ones, we introduced minor changes to the procedure. In essence, a series of images of CB-stained sections covering an entire brain hemisphere (from 146 to 156 sections, depending on the case) were downsampled to a resolution of 15 µm per pixel.

Next, the parts of the image representing brain tissue of a single hemisphere were selected, while the remaining voxels (contralateral hemisphere and the cerebellum) were discarded. The masking procedure was conducted using open-source ITK-SNAP 3.8.0 application (Yushkevich et al., 2006; http://itksnap.org; RRID:SCR_002010). Subsequently, the sections underwent a series of two-dimensional affine alignments to each other and to corresponding template cross-sections to obtain a rudimentary reconstruction that matches the template brain hemisphere and accounts for deviations from the exact coronal sectioning plane. The affine alignment was followed by deformable corrections to refine the reconstruction and make it more suitable for the mapping into the reference template (Fig. S2B).

The registration was carried out by the Advanced Normalization Tools (ANTs) software suite (Avants et al., 2011; http://stnava.github.io/ANTs/; RRID:SCR_004757) with parameters akin to those specified in Majka et al., (2016) and Majka et al., (2020). The process was simultaneously driven by a cross-correlation image similarity metric (Avants et al., 2008) and a set of label maps (Fig. S2B). The label maps were cortical areas outlined manually on Nissl sections (adjacent to corresponding CB-stained sections, Fig. S2A) by an expert (M.G.P.R), based on both cyto- and myelo-architecture, and only then transferred onto the CB-stained sections. An analogous set of label maps was defined in the template (Fig. S2B). Subsequently, corresponding labels were forced to match during registration, which improved the reconstruction’s convergence, speed, overall accuracy, and helped to mitigate issues related to the cross-modal (CB to Nissl) nature of the mapping.

#### Segmentation into cortical areas and layers

The 3D reconstruction procedure resulted in a spatial mapping between the experimental dataset’s coordinate system and the reference template’s stereotaxic coordinates. We then used these transformations to map the segmentation into cortical areas from the template onto the experimental cases (Fig. S2D, left).

Segmentation of layer 4 on CB-stained sections was carried out manually on every other section in each hemisphere using adjacent Nissl-stained sections as a reference (Fig. S2D, middle). The border between layers 1 and 2 was defined computationally as the steepest increase of the CB^+^ neurons density starting from the pial surface. This resulted in the segmentation of the cortex into supragranular layers (layers 2 and 3), granular cell layer (layer 4), and infragranular layers (layers 5 and 6).

In addition, to account for occasional tissue distortions (tears, folds, ruptures, etc.) and local staining artifacts, a separate mask was introduced to exclude these regions from any quantitative analyses. The mask was defined manually by closely inspecting individual sections for any of the mentioned defects.

#### Whole-brain CB^+^ density maps

The CB^+^ neurons density maps were generated for all sections in all cases. They were then downsampled from the native resolution (0.4974 µm per pixel, ×20 magnification) to a mesoscale resolution of 40 µm per pixel. Subsequently, they underwent spatial transformations computed in previous steps. During this procedure, the density maps were corrected by the Jacobian determinant of affine and deformable transformations to maintain accurate values (i.e. the per-section sum of all CB^+^ neurons on the microscopic resolution maps and the spatially transformed maps are preserved). With the 1) density maps of all CB^+^ neurons, 2) segmentation into cortical areas, and 3) segmentation into layers, it was now possible to calculate the average densities for each area and its laminar divisions (Fig. S1C, left) by computing a voxel-wise average density within a mask created by intersecting relevant areal and laminar segmentations.

We excluded the amygdalopiriform transition area (APir) from the analyses due to its small size and difficulties in identifying the precise boundaries of this area, either manually or by the registration algorithm. In addition, we decided to exclude the entorhinal cortex (Ent), piriform cortex (Pir), area 24a, (A24a), area 29a-c (A29a-c), and area 35 (A35) from any analyses involving laminar divisions due to the lack of a clearly defined layer 4 homolog.

### QUANTIFICATION AND STATISTICAL ANALYSIS

ANOVA analyses presented in Figures 3 and 7 were carried out using the scikit-posthocs Python package (v. 0.7.0) using the Kruskal–Wallis H test. Statistical significance was assessed according to post-hoc Dunn’s test with Holm–Bonferroni correction for multiple comparisons.

## ADDITIONAL RESOURCES

**Supplemental Figure S1.**
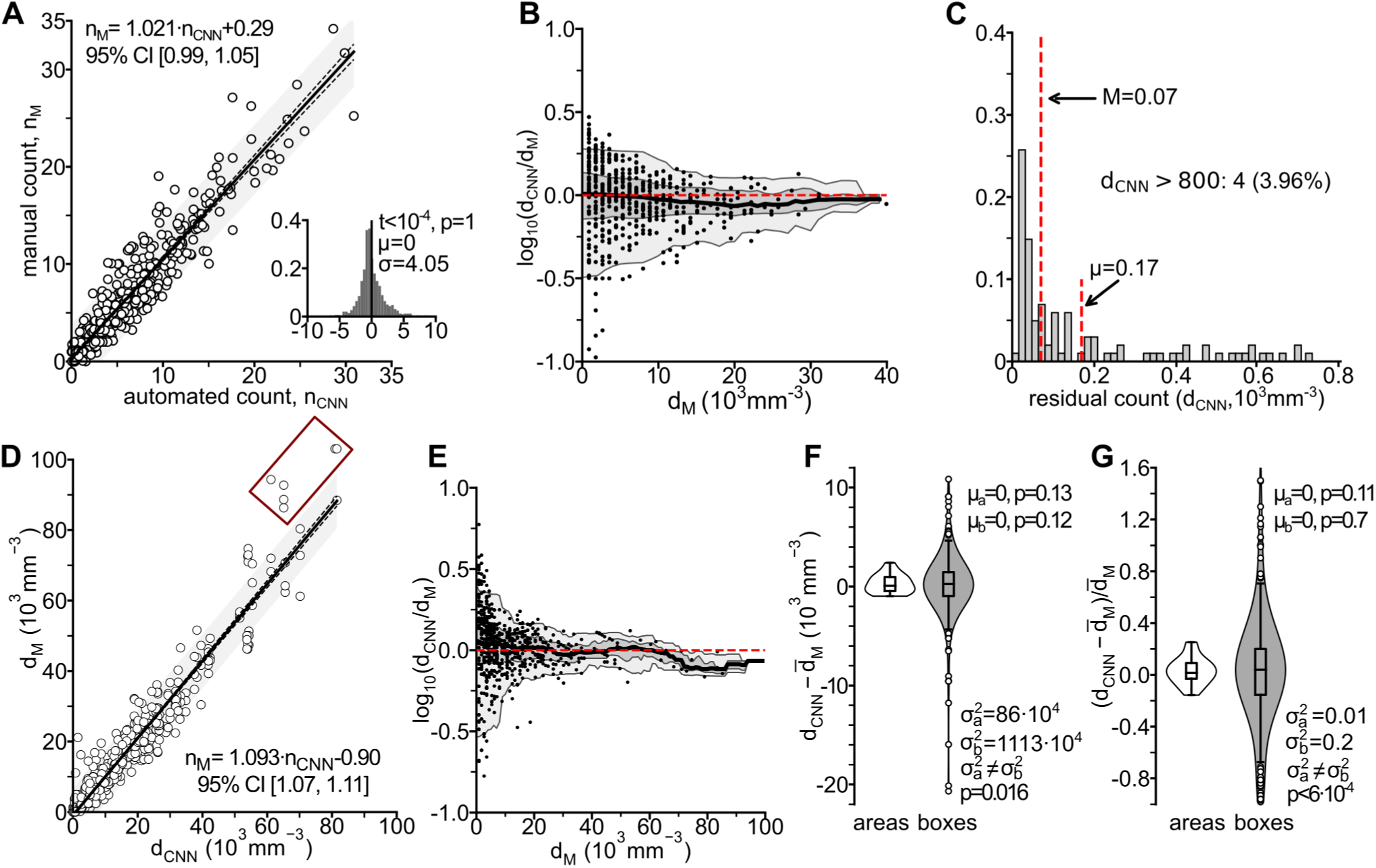
Detailed evaluation of the automated cell counting method. (A) The evaluation of the automatic cell counting performance using the validation dataset (603 counting boxes). The relation between the automated count (n_CNN_, abscissas) and the ground truth manual count (n_M_, ordinates) shows a linear relation between the two quantities. The residuals average to zero (two-sided t-test, t<10^-4^, p=1, µ=0). (B) Relative error of the U-Net CNN (d_CNN_) neuronal densities against those established by manual counting (d_M_). Black points represent results for the individual counting boxes. Order statistics (median: thick black line, the light gray bands: 5^th^ and 95^th^ centiles, dark gray: lower and upper quartiles) calculated locally within a 5·10^3^mm^-3^ wide moving window. The red dashed line represents the agreement between the U-Net CNN and the manual results. The relative error decreases with increasing CB^+^ densities (the bands are wider for lower densities and narrower for higher ones). (C) Histogram of the residual densities (i.e., the density estimated by the U-Net CNN within image patches known to contain no neurons). Among the 2,216 empty image patches, 50% had a residual density less or equal to 70 mm^-3^, and the mean residual density amounted to 170 mm^-3^, which shows that the impact of the background on the U-Net CNN results is negligible. (D) Comparison of the CB^+^ densities estimated by the U-Net CNN against three expert neuroanatomists. Densities established by the U-Net CNN (d_CNN_, abscissas) against the count obtained manually (d_M_, ordinates) for each image patch in these areas (939 values in total) show a linear relation between the two quantities. (E) Relative error of the U-Net CNN (d_CNN_) neuronal densities against manual counting (d_M_) for the benchmark dataset. Black points represent results for the individual image patches. Order statistics (median: thick black line, light gray bands: 5^th^ and 95^th^ centiles, dark gray: lower and upper quartiles) calculated locally within a 5·10^3^ mm^-3^ wide moving window. The dashed red line represents the agreement between the U-Net CNN and the manual results. The densities based on U-Net CNN match those established by manual plotting for all counting boxes except those with the highest densities (>70·10^3^ mm^-3^), such as those obtained in the granular layer of V1 and V2. For these, the U-Net CNN densities tend to be lower than those established manually, likely reflecting occurrences of multiple, superimposed small cells. (F, G) Analyses of the discrepancies between the d_CNN_ and the *d̅*_M_ densities, either for absolute (E) or normalized (i.e. divided by the average of the three expert observers, *d̅*_M_) (F) density values. In either approach, the mean difference is statistically indifferent from zero. However, per-box comparisons (gray violin plots) exhibit higher variance than the per-area comparisons (white violin plots).

**Supplemental Figure S2.**
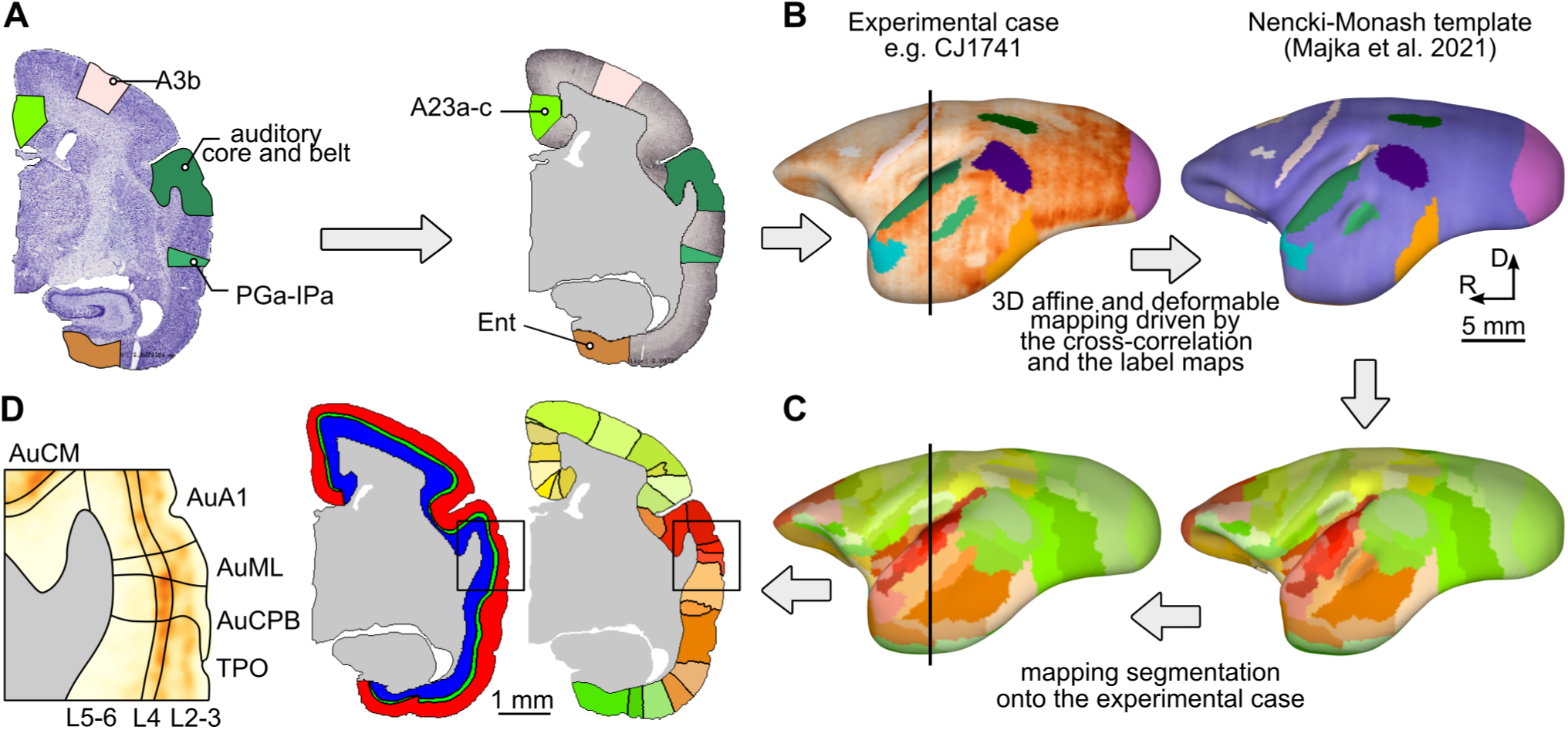
Cytoarchitecture-aware registration to Nencki-Monash reference template (Majka et al., 2021). (A) Label maps outlined manually on Nissl sections (*left*) are transferred onto the adjacent CB-stained sections (*right*). (B) Registration of the experimental case to the reference template. A combined view of the CB-stained sections and the outlined label maps (*left*) in the experimental case, the Nissl-stained section, and the label maps in the reference template (*right*). (C) The computed spatial transformations are used to map the segmentation from the reference template (*right*) onto the experimental case (*left*). The black line indicates the coronal location of the section shown in panels A and D. (D) Example CB-stained coronal section segmented into cortical areas (*right*), manual segmentation of the same section into supragranular, granular, and infragranular layers (red, green and blue, respectively, *middle*). The combination of the areal and laminar segmentation allows for computing densities of CB^+^ neurons in individual areas across different layers (*left*). See Supplemental Table 1 for a list of areas and their abbreviations.

## Notes

https://www.marmosetbrain.org/whole_brain_cb_maps

### Competing Interest Statement

The authors have declared no competing interest.

